# CRISPR screen reveals that EHEC’s T3SS and Shiga toxin rely on shared host factors for infection

**DOI:** 10.1101/316919

**Authors:** Alline R. Pacheco, Jacob E. Lazarus, Brandon Sit, Stefanie Schmieder, Wayne I. Lencer, Carlos J. Blondel, John G. Doench, Brigid M. Davis, Matthew K. Waldor

**Affiliations:** Division of Infectious Diseases, Brigham and Women’s Hospital. Boston, MA, USA; Department of Microbiology and Immunobiology, Harvard Medical School. Boston, MA, USA; Division of Infectious Diseases, Massachusetts General Hospital. Boston, MA, USA; Division of Gastroenterology, Boston Children’s Hospital. Boston, MA, USA; Department of Pediatrics, Harvard Medical School. Boston, MA, USA; Department of Pediatrics, Harvard Digestive Diseases Center. Boston, MA, USA; Institute of Biomedical Sciences, Universidad Autónoma de Chile. Santiago, Chile; Broad Institute of MIT and Harvard. Cambridge, MA, USA; Howard Hughes Medical Institute, Boston, MA, USA

**Keywords:** CRISPR screen, host susceptibility, T3SS, Shiga toxin, EHEC, EPEC, sphingolipid synthesis, TM9SF2, LAPTM4A

## Abstract

Enterohemorrhagic *Escherichia coli* (EHEC) has two critical virulence factors – a type III secretion system (T3SS) and Shiga toxins (Stx) – that are required for the pathogen to colonize the intestine and cause diarrheal disease. Here, we carried out a genome-wide CRISPR/Cas9 loss-of-function screen to identify host loci that facilitate EHEC infection of intestinal epithelial cells. Many of the guide RNAs identified targeted loci known to be associated with sphingolipid biosynthesis, particularly for production of globotriaosylceramide (Gb3), the Stx receptor. Two loci (TM9SF2 and LAPTM4A) with largely unknown functions were also targeted. Mutations in these loci not only rescued cells from Stx-mediated cell death, but also prevented cytotoxicity associated with the EHEC T3SS. These mutations interfered with early events associated with T3SS and Stx pathogenicity, markedly reducing entry of T3SS effectors into host cells and binding of Stx. The convergence of Stx and T3SS onto overlapping host targets provides guidance for design of new host-directed therapeutic agents to counter EHEC infection.

**Importance:** Enterohemorrhagic *Escherichia coli* (EHEC) has two critical virulence factors – a type III secretion system (T3SS) and Shiga toxins (Stx) – that are required for colonizing the intestine and causing diarrheal disease. We screened a genome-wide collection of CRISPR mutants derived from intestinal epithelial cells and identified mutants with enhanced survival following EHEC infection. Many had mutations that disrupted synthesis of a subset of lipids (sphingolipids) that includes the Stx receptor globotriaosylceramide (Gb3), and hence protect against Stx intoxication. Unexpectedly, we found that sphingolipids also mediate early events associated with T3SS pathogenicity. Since antibiotics are contraindicated for the treatment of EHEC, therapeutics targeting sphingolipid biosynthesis are a promising alternative, as they could provide protection against both of the pathogen’s key virulence factors.

## Introduction

Enterohemorrhagic *E. coli* (EHEC) is a food-borne human pathogen that causes diarrheal illness worldwide. Infection is often associated with bloody diarrhea that is usually self-limited; however, 5-7% of cases progress to hemolytic uremic syndrome (HUS), a life-threatening complication that can result in renal failure and neurological sequelae (1). EHEC pathogenesis shares many features with that of enteropathogenic *E. coli* (EPEC), another extracellular pathogen that colonizes the intestine. Successful colonization by both species is dependent upon a type III secretion system (T3SS) that enables tight adherence of bacteria to host epithelial cells by inducing characteristic actin cytoskeletal rearrangements and loss of microvillus structure (attaching and effacing (AE) lesions) (2). EHEC virulence is also markedly shaped by production of Shiga toxins (Stx), variants of which are often present in multiple copies within the EHEC genome. Translocation of Stx to tissues outside of the intestinal tract is thought to underlie the development of HUS (3, 4).

The EHEC T3SS injects a plethora of effector proteins into host cells, resulting in alteration or disruption of numerous host cell processes. During infection, EHEC is thought to target epithelial cells within the large intestine; however, a variety of cultured cell lines have been used to characterize the activity of this system. In vivo and in vitro studies have revealed that a key effector is the Translocated Intimin Receptor (Tir) (5). Tir is inserted into the host cell membrane and serves as a receptor for the bacterial adhesin, intimin (6). Interactions between intimin and Tir are also required for recruitment and rearrangement of actin and other cytoskeletal proteins underneath adherent bacteria, which results in characteristic actin-rich “pedestals.” In animal models, deletions of *tir* or *eae* (the intimin locus) and mutations that render the T3SS inactive markedly reduce the pathogen’s capacity to colonize the intestine and cause disease (7, 8).

Thirty eight bacterial proteins in addition to Tir have been confirmed as type 3 secreted effector proteins in EHEC (9). Unlike structural components of the T3SS, individual effector proteins are frequently not essential for bacterial virulence; although their roles have not been fully defined, it is clear that effector proteins can act in redundant, synergistic and antagonistic fashions (10). Key host processes modulated by EHEC effectors include innate immunity, cytoskeletal dynamics, host cell signaling, and apoptosis (11). EHEC effectors also restrict host cell phagocytosis of this extracellular pathogen. Effectors undergo an ordered translocation and after its translocation, the effector protein EspZ functions as a “translocation stop” that prevents unlimited effector translocation and reduces infection-associated cytotoxicity (12). Compared to wt infection, in vitro infection with *espZ*-deficient strains results in greater host cell detachment, loss of membrane potential, and formation of condensed nuclei (13).

Although Stx are pivotal to EHEC pathogenesis, the effects of these AB_5_ toxins on the intestinal epithelium per se are not entirely clear. Toxicity was initially thought to be largely restricted to tissues beyond the intestinal tract (e.g., microvascular endothelial cells within the kidneys and the brain in the setting of HUS) (14); however, more recent in vivo and ex vivo studies suggest that Stx intoxication may also occur in the intestine at the primary site of infection. Although at low levels, receptors for Stx are present within human colonic epithelial cells (15), and Stx2 causes extensive cell death to the intestinal mucosa (16, 17). Furthermore, oral administration of Stx can lead to diarrhea in animals, and in several animal models of EHEC intestinal disease, severe diarrhea is dependent on Stx (8, 17).

The principal receptor for most forms of Stx (including Stx1 and Stx2, which are produced by the EHEC strain used in this study) is a neutral glycosphingolipid, globotriaosylceramide (Gb3). Following binding of Stx to Gb3, the toxin is internalized and undergoes retrograde transport through early endosomes, the Golgi, and the ER; the A subunit is cleaved by furin in the Golgi, followed by disulfide bond reduction in the ER that releases the catalytic active A1 fragment, which undergoes retro-translocation into the cytosol (18). Site specific depurination of 28S rRNA by the toxin results in inhibition of protein synthesis and can induce the ribotoxic stress response, the unfolded protein response, and apoptosis (19–22).

Analyses of EHEC pathogenesis have primarily focused upon identification and characterization of bacterial factors rather than on host factors required for pathogenicity. Though some host factors, particularly those required for the actions of Stx and of the T3SS effectors, have been identified, to date, unbiased genome-wide screens for EHEC susceptibility loci have not been reported. Recently, such screens have become possible, given the advent of CRISPR/Cas9-based libraries of host mutants whose composition can be monitored using high throughput DNA sequencing. We recently used this approach to screen for host factors that mediate susceptibility to *V. parahaemolyticus’* two T3SS, and identified several host processes not previously linked to T3SS activity (23). The efficiency and power of this approach (24, 25), prompted us to adopt this approach to identify mutants with heightened resistance to EHEC.

Here, we identify and characterize intestinal epithelial cell mutants that become enriched following library infection with an EHEC strain producing an active T3SS as well as Stx. Although minimal overlap between the action of the T3SS and Stx have previously been reported, we identified several host loci and processes that are required for the effects of both virulence factors. Genes required for production of the Stx receptor Gb3 and other sphingolipids were also found to be necessary for translocation of T3SS effectors into host cells. Additionally, we identified 2 minimally characterized loci not previously linked to either T3SS or Stx response pathways and find that they are critical for the biogenesis of host cell Gb3 and hence susceptibility to EHEC infection.

## Results

### CRISPR/Cas9 screen for host factors conferring susceptibility to EHEC infection

We developed a genome-wide CRISPR/Cas9 screen to identify host factors that contribute to susceptibility to EHEC infection in the HT-29 colonic epithelial cell line using the Avana library of sgRNAs (23). This library contains four sgRNAs targeting each of the annotated human protein coding genes (26). Host cells were infected with a Δ*espZ* derivative of EHEC strain EDL933 (which carries genes encoding both Stx1 and Stx2). The Δ*espZ* mutation heightens T3SS activity and increases host cell death associated with infection by EHEC (13) (Fig. S1A). We anticipated that if we could increase the toxicity of the EHEC T3SS (Fig. S1B), we would enhance the screen’s selective pressure and yield greater enrichment of host cell mutants resistant to this key virulence system. Although Stx1 and Stx2 were also produced under infection conditions (Fig. S1C), initial toxicity assays using purified toxin suggested that they would not exert substantial selective pressure during the screen. In contrast to infection of cells with Δ*espZ* EHEC, which resulted in marked (~80%) host cell death by the end of the infection period (Fig. 1AB), a corresponding 6-hour treatment of cells with purified toxin exceeding the amount detected during infection had minimal effect on viability (Fig. S1D).

**Figure 1.**
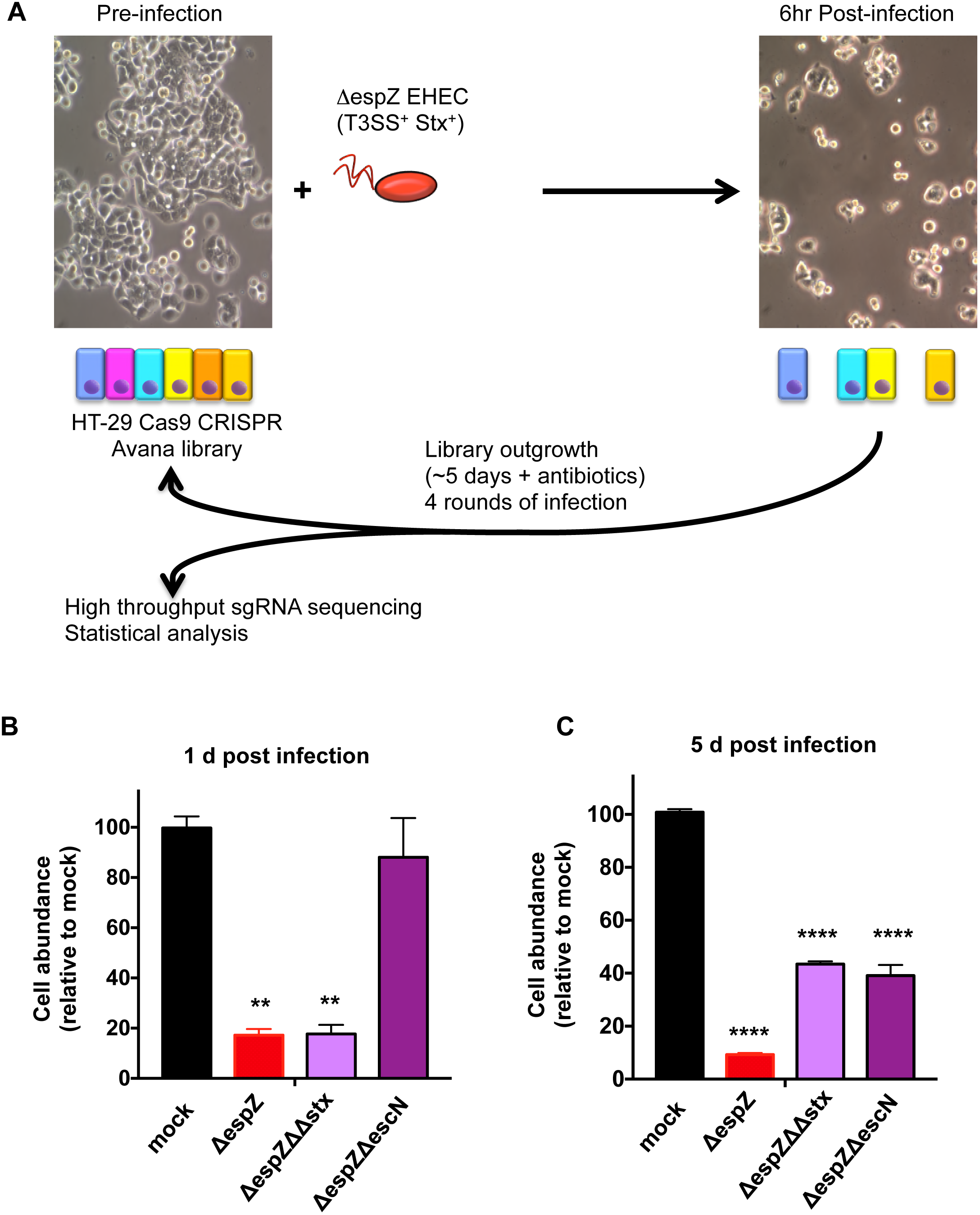
Design of a CRISPR/Cas9 screen to identify host factors underlying susceptibility to EHEC infection. A) Schematic of the infection and outgrowth process for an HT-29 Cas9/CRISPR library undergoing multiple rounds of infection with Δ*espZ* EHEC, which has an active T3SS and secretes Stx1 and Stx2. B, C) Abundance of HT29 cells infected with the indicated strain relative to the abundance of mock-infected cells at day 1 (B) and day 5 (C) post-infection. Graphs display mean and SD from 3 independent experiments. P values (**P<0.01, ****P<0.0001)

For the screen, two biological replicates of libraries of HT-29 cells mutagenized with the Avana guide RNAs were infected for 6 hr with Δ*espZ* EHEC at an MOI of 100 (Fig. 1A). Following infection, resistant cells were cultured in the presence of antibiotics until reaching ~70% confluency (~5 days), then reseeded and reinfected. Genomic DNA was isolated from a fraction of the surviving population after each of the four rounds of infection as well as from the initial uninfected cells, and high throughput sequencing of integrated sgRNA templates was performed to indirectly quantify the abundance of the associated mutants. We found that as our screen progressed, as would be expected for a library under strong selection, representation became biased toward a subset of enriched genes (Fig. S2A). Statistical analysis was performed using the STARS algorithm, which integrates data from independent guides targeting the same gene to identify the most enriched genotypes (26).

We identified 13 loci with statistically significant enrichment (p< 0.001) in both libraries after 4 rounds of infection (Fig. 2A). Unexpectedly, given our results with purified toxin, more than half of the enriched loci encoded factors associated with sphingolipid biosynthesis, and many were closely connected to synthesis of Gb3, the Stx receptor (Fig. 2B). For example, hits included the Golgi-localized enzymes A4GALT, which catalyzes the final step in Gb3 synthesis; B4GALT5, which catalyzes production of the A4GALT substrate lactosylceramide; and UGCG, which converts ceramide (a precursor for all glycosphingolipids) into glucosylceramide. Additional hits included the ER-localized SPTLC2, SPTSSA, and KDSR, all of which lie on the ceramide synthesis pathway, and ARF1, which indirectly regulates intracellular trafficking of glucosylceramide (27).

**Figure 2.**
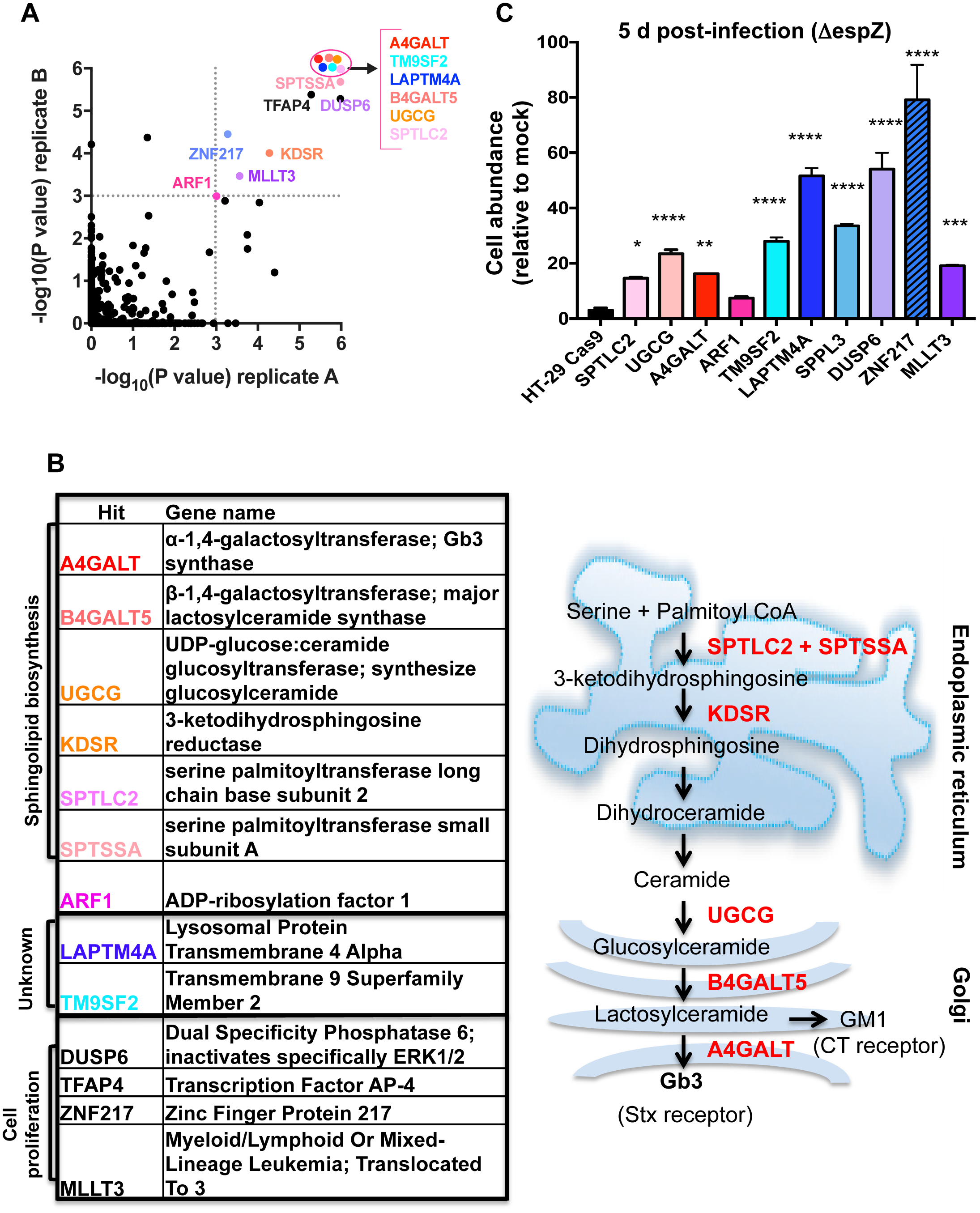
Mutations that disrupt sphingolipid biosynthesis and poorly characterized genes are enriched in the HT-29 CRISPR/Cas9 library following repeated infection with *espZ* EHEC. A) Scatterplot of the statistical significance in each library (A and B) associated with the genes ranked in the top 5% by the STARS algorithm. Genes with a p value <= 0.001 in both libraries (upper right quadrant) are named; genes within the ellipse all have p values <2.0e^-06^. B) Products of genes shown in (A) with p<=0.001 in both libraries and schematic representation depicting the subcellular localization of enzymes (black) that contribute to sphingolipid biosynthesis. A subset of substrates/products are depicted in red. C) Abundance of HT29 control and mutant cells infected with Δ*espZ* EHEC relative to the abundance of mock-infected cells at day 5 post infection. Graphs display mean and SD from 3 independent experiments compared to HT-29 Cas9 (leftmost bar). P values (* P<0.05, ** P<0.005, *** P<0.001, ****P<0.0001)

Enriched loci also included two genes, TM9SF2 and LAPTM4A, whose functions are largely unknown. Although both are members of larger gene families of structurally related proteins, only guide RNAs targeting these particular family members were found to be enriched, suggesting that they have specific functions conferring susceptibility to EHEC infection (Fig. S2B). Human TM9SF2 has been reported to be a Golgi-resident transmembrane protein required for the Golgi localization of NDST1, a sulfotransferase (28). Its homologs in Drosophila (TM9SF2/4) and Dictyostelium (Phg1A/Phg1C) have been linked to innate immunity and membrane protein localization (29, 30). LAPTM4A has been reported to encode a transmembrane protein localized to lysosomes and late endosomes. It has been linked to intracellular transport of nucleosides, multidrug resistance, and maintenance of lysosomal integrity (31–34). In addition to the enrichment of guides targeting TM9SF2, LAPTM4A, and genes related to sphingolipid biosynthesis, our analysis detected enrichment for guides targeting several genes associated with cancer and cell proliferation (MLLT3, TFAP4, ZNF217, and DUSP6) (35–38).

The prominence among our hits of Gb3-related genes was unanticipated, because our preliminary studies suggested that T3SS rather than Stx would exert the strongest selective pressure in our screen. T3SS from several organisms have been hypothesized to associate with lipid rafts (39–41), transient membrane microdomains which typically are enriched in sphingolipids (including Gb3) (19); however, studies of T3SS activity and host membrane components have generally focused on the importance of cholesterol, and a role for Gb3 in EHEC pathogenesis beyond that of Stx receptor has not been reported. To more precisely define the contribution of screen hits to susceptibility to EHEC infection, we developed assays that enabled the effects of Stx and T3SS on HT-29 cells to be investigated independently. Host cells were infected in parallel with an Δ*espZ*, an Δ*espZ* Δ*escN* mutant (which lacks an ATPase essential for T3SS activity), an Δ*espZ* Δ*stx1* Δ*stx2* mutant (ΔΔ*stx*, which does not produce Stx1 or Stx2), or mock infected, and the number of host cells present after 1 or 5 days of infection was determined (Fig. S1B). These experiments revealed similar marked declines in abundance of HT-29 cells 1 day post-infection with the type 3-active Δ*espZ* or Δ*espZ* ΔΔstx EHEC (to <20% of that seen in mock infected cells; Fig. 1B); in contrast the type 3-deficient but toxin-producing Δ*espZ* Δ*escN* infection had no significant effect on host cell abundance at this time point (Fig. 1B). However, in the subsequent 4 days, there was a marked difference in growth between cells infected with the Δ*espZ* vs. Δ*espZ* ΔΔ*stx* strains. Population expansion of cells previously exposed to toxin (Δ*espZ* infection) was far slower than that of cells that were never exposed to the toxin (Δ*espZ* ΔΔ*stx* infection) (Fig. 1C, Fig. S1B). The abundance of HT-29 cells infected with the Δ*espZ*Δ *escN* strain also differed significantly from that of mock infected cells by day 5, likely further reflecting the consequence of toxin exposure at this time point. Collectively, these analyses demonstrate that the impact of T3SS is most clearly evident 1 day post infection, and can be differentiated from that of toxin via infection with the Δ*espZ* ΔΔ*stx* strain. In contrast, the effects of Stx are delayed and become more apparent 5 days post infection, and can be assayed in infection with the T3SS-deficient Δ*espZ* Δ*escN* strain. Given that our screen included multiple rounds of 5-day outgrowth following infection (during which effects of Stx could manifest), we conclude that the screen enriched for mutants resistant to Stx as well as T3SS.

For initial validation of our screen hits, mutants corresponding to selected enriched loci were constructed and verified by Sanger sequencing or Western blot (Fig. S2C-E), and their abundance (relative to mock infected cells) was assessed 5 days post-infection with Δ*espZ* EHEC. In comparison to the HT-29 Cas9 control strain, all but one mutant (ARF1) had significantly enhanced abundance at 5 days post-infection (Fig. 2C), suggesting that our selection and analysis yielded robust and reliable data regarding susceptibility to EHEC infection.

### Sphingolipid biosynthesis facilitates T3SS killing

Because the hits predicted to be involved in cell proliferation (i.e. DUSP6, TFAP4, ZNF217 and MLLT3) were less likely to be involved in EHEC pathogenesis per se, we chose to focus further studies on a subset of the sphingolipid biosynthesis mutants and on mutants in the largely uncharacterized loci TM9SF2 and LAPTM4A. In particular, we focused on factors mediating synthesis of glycosphingolipids, particularly A4GALT, which should only have impaired production of Gb3 and other globo-series glycosphingolipids (42), and UGCG, which is required for synthesis of all glycosphingolipids except galactosylceramines (43). ARF1 was also included, due to its contribution to intracellular trafficking of UGCG’s product from the cis-medial Golgi to the trans Golgi network, where B4GALT5 and A4GALT are found (44). To assess the bacterial factors underlying the enrichment of these loci, mutants were first infected with the Δ*espZ* ΔΔ*stx* strain or mock infected, and cell abundance was assessed one day following infection. Notably, all of the mutant cells displayed significantly elevated relative abundance compared to wt HT-29 cells (Fig. 3A). Coupled with our prior analyses of bacterial factors modulating host survival at this time point (Fig. 1B), these results suggest that these mutations confer resistance to Δ*espZ* ΔΔ*stx* infection by protecting against the effects of EHEC’s T3SS, and thus that associated loci may play a role in the host cell response to T3SS. Of these factors, only ARF1 has previously been linked to T3SS-mediated processes; it is thought to facilitate insertion of the T3SS translocon during Yersinia infection (45) but has not been linked to T3SS activity in EHEC.

**Figure 3.**
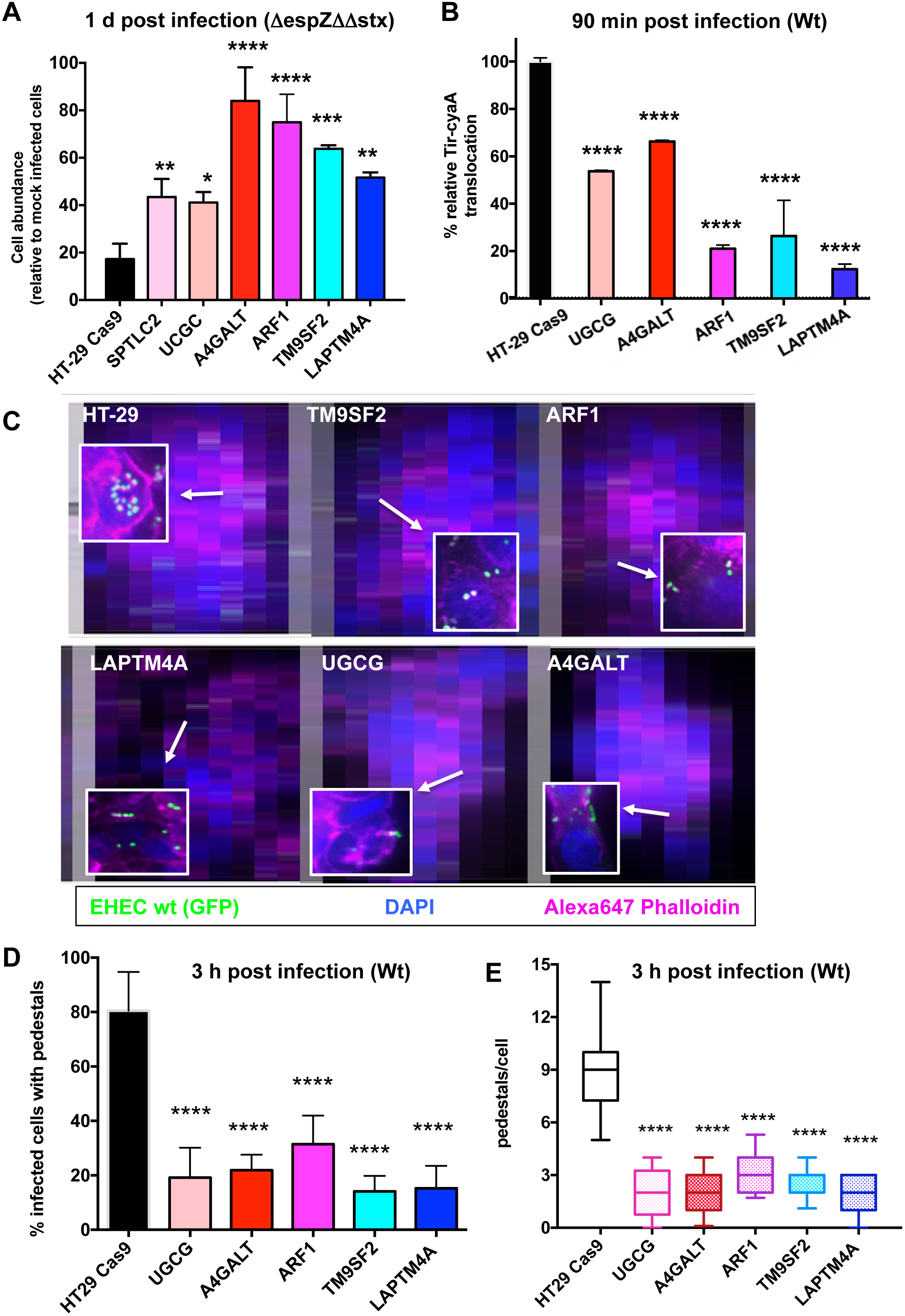
Disruption of host sphingolipid biosynthesis genes and poorly characterized genes reduces the activity and cytotoxicity of EHEC’s T3SS. A) Abundance of control and mutant HT29 Cas9 cells infected with Δ*espZ* Δ*stx1* Δ*stx2* EHEC relative to the abundance of mock-infected cells at day 1 post infection. Graphs display mean and SD from 3 independent experiments. P values (* P<0.02, ** P<0.01, *** P < 0.001, **** P<0.0001) were obtained from one-way ANOVA with Dunnet post-correction. B) Relative translocation of Tir-CyA from wt EHEC into HT-29 Cas9 control cells and the indicated HT-29 mutants, based on cAMP levels. Translocation into HT-29 Cas9 control cells was set as 100%. Data reflect mean and SD from 3 independent experiments. P values (**** P<0.0001) are based on one-way ANOVA with Dunnet post-correction. C) Confocal microscopy of control and mutant HT-29 Cas9 cells infected for 6 hr with GFP-EHEC, then stained for F-actin with Alexa647-phalloidin (pink) and DAPI (blue; labels nuclei). Merged images are shown. Focal colocalization of bacteria and actin reflects formation of actin pedestals. White boxes show enlarged image, to highlight pedestals. D) Percentage of the indicated host cells with actin pedestals 6 hr after infection. 250 cells were assessed for each host genotype. E) Number of pedestals per host cell; box plots show range (min to max) of pedestal numbers. 100 cells with AE lesions were counted per genotype. P values (**** P<0.0001)

The first effector translocated by EHEC’s T3SS – Tir – is essential for the activity of this secretion system. In the absence of Tir, translocation of additional effectors does not occur, nor does the characteristic cytoskeletal rearrangement and formation of membrane “pedestals” underneath adherent bacteria (5, 46). Tir translocation can be assessed using a Tir-CyaA reporter fusion protein, followed by measurement of intracellular cAMP levels (47). We monitored translocation of this reporter, which is dependent on an intact T3SS, from wt EHEC into control HT-29 Cas9 cells and mutants that appeared resistant to the effects of T3SS. These experiments revealed significantly lower Tir translocation into all mutants than into control HT-29 cells, ranging from ~70% (A4GALT) down to ~10% (LAPTM4A) of wt levels (Fig. 3B). Consistent with this observation, immunofluorescence microscopy of wt and mutant HT-29 cells infected with wt EHEC revealed markedly fewer adherent bacteria and associated actin-rich pedestals. While EHEC formed pedestals on ~80% of infected wt HT-29 cells, pedestals were detected on only 18-35% of infected mutants tested, and fewer pedestals were generally observed per mutant cell (Fig. 3CDE, Fig. S3A). Collectively, these results indicate that the mutations rendering HT-29 cells less susceptible to the cytotoxic effects of EHECs T3SS all limit early steps in the T3SS effector translocation process, although they do not fully disrupt this process. These results also demonstrate that the mutants identified in our screen are protective against EHEC even when its T3SS activity has not been augmented by mutation of Δ*espZ*.

EHEC’s T3SS machinery and associated effectors are similar to those of enteropathogenic *E. coli* (EPEC), a related pathogen that does not produce Shiga toxin but that also requires its T3SS to colonize and cause disease in the human intestine (48). We investigated whether the mutations that protect HT-29 cells from EHEC infection also render HT-29 less susceptible to EPEC. Wt and mutant HT-29 cells were infected with an EPEC Δ*espZ* mutant that, like its EHEC counterpart, is reported to have increased cytotoxicity relative to the wt strain (13, 49) (Fig. S3B). All 5 mutants tested exhibited increased survival compared to the wt cells at 1 day post infection, and this increase was linked to the presence of a functional T3SS (Fig. S3C). Survival of the TM9SF2 and ARF1 mutants was particularly enhanced, with the number of infected cells nearing 50% of the mock infected controls, compared to the ~5% observed with wt HT-29. Overall, our observations suggest that host glycosphingolipids modulate T3SS-mediated cytotoxicity for both pathogens, although pathogen reliance on particular sphingolipids is not necessarily conserved. For example, the absence of A4GALT had a dramatic effect on EHEC cytotoxicity, but only a modest influence on EPEC cytotoxicity. Additionally, our results suggest that TM9SF2 may play a conserved role in facilitating the activity of the EHEC and EPEC T3SS.

Given previous reports that lipid rafts may promote T3SS activity, that Gb3 high-density association within lipid rafts is important for Stx binding (50), and our screen’s identification of numerous sphingolipid-related loci, we hypothesized that TM9SF2 and LAPTM4A mutants might be less susceptible to EHEC infection due to alterations in lipid raft production or dynamics. To explore these possibilities, we compared the trafficking in control, TM9SF2 and LAPTM4A cells of a chimeric GPI-anchored GFP construct (GPI-GFP), which is transported to the plasma membrane where it becomes enriched in lipid rafts (51). Cell surface fluorescence was similarly homogenous in all three genetic backgrounds, and we could not detect consistent differences in steady state plasma membrane fluorescence between the 3 samples (Fig. S3D). To determine if kinetics of trafficking and insertion might nevertheless differ between wt and mutant cells, we performed quantitative photobleaching. The rate of signal decay was similar for all 3 backgrounds, suggesting that bulk plasma membrane trafficking and lipid raft insertion is not grossly disrupted in TM9SF2 and LAPTM4A cells (Fig. S3E). Further studies will be needed to define the precise means by which these mutations, as well as others tested above, limit susceptibility to EHEC and EPEC’s T3SS.

### LAPTM4A and TM9SF2 are required for Gb3 biosynthesis

As noted above, the effects of Stx on HT-29 abundance were evident 5 days post-infection, and could be clearly distinguished from those of EHEC’s T3SS through infection with the Stx+ T3SS-deficient Δ*escN* mutant (Fig. 1C). Therefore, we also compared the abundance of wt HT-29 cells and several mutants from our panel after challenge with the Δ*escN* strain. As anticipated, mutants lacking the sphingolipid biosynthesis factors A4GALT, UGCG, and SPTLC2 (all of which contribute to Gb3 production) were far less susceptible to Δ*escN* infection than wt HT-29 cells; at 5 days post infection, the abundance of these mutants did not differ from that of mock-infected controls (Fig. 4A). Intriguingly, the TM9SF2 and LAPTM4A mutants were also significantly more abundant than wt cells by 5 days post Δ*escN* infection, suggesting that these factors not only contribute to resistance to T3SS-mediated cytotoxicity, but also could be host factors facilitating intoxication.

**Figure 4.**
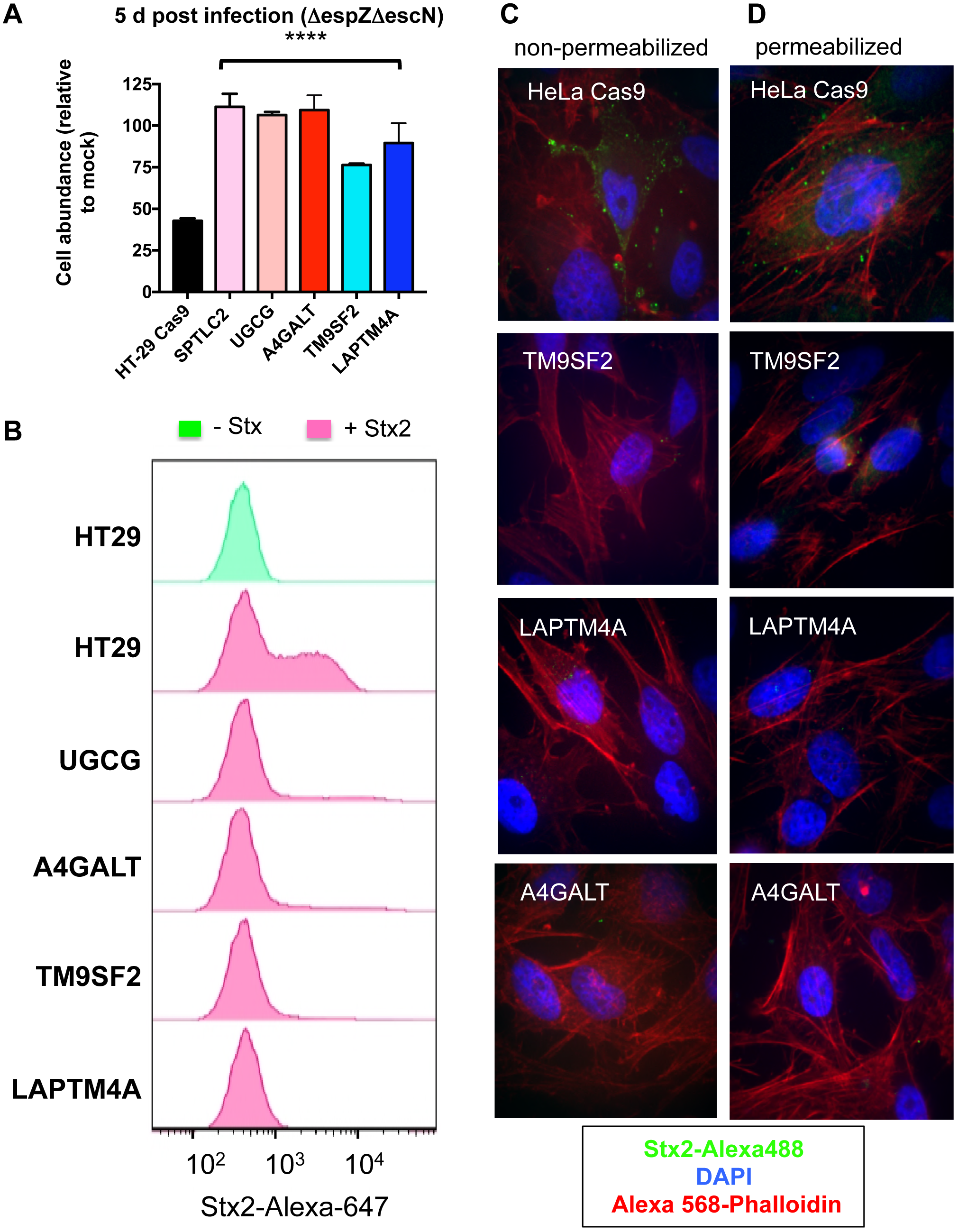
TM9SF2 and LAPTM4A promote sensitivity to Stx. A) Abundance of wt and mutant HT29 Cas9 cells infected with T3SS-deficient EHEC (Δ*espZ* Δ*escN)* relative to the abundance of mock-infected cells at day 5 post infection. P values (**** P<0.0001) are based on one-way ANOVA with Dunnet post-test correction. B) Flow cytometry analysis of Stx2-Alexa647 binding to wt and mutant HT-29 Cas9 cells. Histograms show the distribution of fluorescence intensity in the total cell population in the presence and absence of toxin. C, D) Confocal microscopy of Stx2-Alexa488 (green) binding to non-permeabilized (C) and permeabilized (D) control and mutant HeLa Cas9 cells. Cells were also stained with DAPI and Alexa568-phalloidin

To begin to explore the means by which TM9SF2 and LAPTM4A mutations protect against Stx, we tested the capacity of mutants to bind to fluorescently tagged toxin. As controls, we also assayed A4GALT and UGCG mutants, which are known to be completely deficient in Gb3 production, and hence cannot bind Stx. Flow cytometry analyses, which were performed both in HT-29 (Fig. 4B) and HeLa (Fig. S4A) cells, revealed that there was a high (but not uniform) level of binding of Stx2 to both wt cell types. Notably, there was marked reduction in Stx binding in both the TM9SF2 and LAPTM4A mutant cells (Fig. 4B and Fig. S4A); these mutants bound equal to or less toxin than the A4GALT and UGCC mutants. The residual binding observed in all genetic backgrounds may reflect non-specific toxin adsorption or low level genetic heterogeneity in the CRISPR/Cas9-mutagenized lines.

Similarly, fluorescence microscopy, which was performed using wt HeLa cells and their derivatives due to their favorable imaging characteristics, did not reveal Stx binding to any of the mutants, even when cells were permeabilized to enable binding to intracellular receptor (Fig. 4C, D). Thus, the TM9SF2 and LAPTM4A mutants’ deficiencies in Stx binding do not appear to reflect impaired trafficking of Gb3 to the cell surface, but instead reflect defective synthesis and/or enhanced degradation of this glycosphingolipid.

To evaluate the specificity of the deficiency in the TM9SF2 and LAPTM4A HT-29 mutants, we used flow cytometry to measure their capacity to bind cholera toxin (CT), which interacts with the glycosphingolipid GM1 (Fig. 2B) (52). In contrast to the near ablation of Stx binding in these mutants, there was a comparatively modest reduction in the binding of fluorescently labeled CT to these cells compared to wt HT-29 cells and A4GALT mutant cells (which produce normal amounts of GM1) (Fig. S4B). As expected, the UGCG cells showed a far more marked decrease in binding to cholera toxin, since UGCG is required for synthesis of GM1 (43). Collectively, these observations suggest that the TM9SF2 and LAPTM4A mutants’ deficiencies in Stx binding reflect relatively specific reductions in production of Gb3 or related globo-series glycosphingolipids, rather than deficiencies that consistently impair synthesis or trafficking to the cell membrane of multiple surface receptors. Coupled with our analysis of the set of mutants that display reduced sensitivity to T3SS-mediated cytotoxicity, these observations suggest that reduced production of Gb3 likely contributes to the TM9SF2 and LAPTM4A mutants’ resistance to the effects of EHEC’s T3SS as well as to Stx.

To gain greater insight into the mechanism by which TM9SF2 and LAPTM4A enable infection by EHEC, we determined their subcellular localization in HeLa cells via confocal microscopy. Consistent with previous reports (28), we found that TM9SF2 co-localized with GM130, a Golgi matrix protein (Fig. 5A). A subset of fluorescence emanated from nucleoli, potentially resulting from non-specific primary antibody binding, though Stx has been reported to be actively transported into nucleoli (53). Unexpectedly, as prior studies localized LAPTM4A to lysosomes (31-34), we found that a LAPTM4A-GFP fusion protein localized to the Golgi, like TM9SF2 (Fig. 5A).

**Figure 5.**
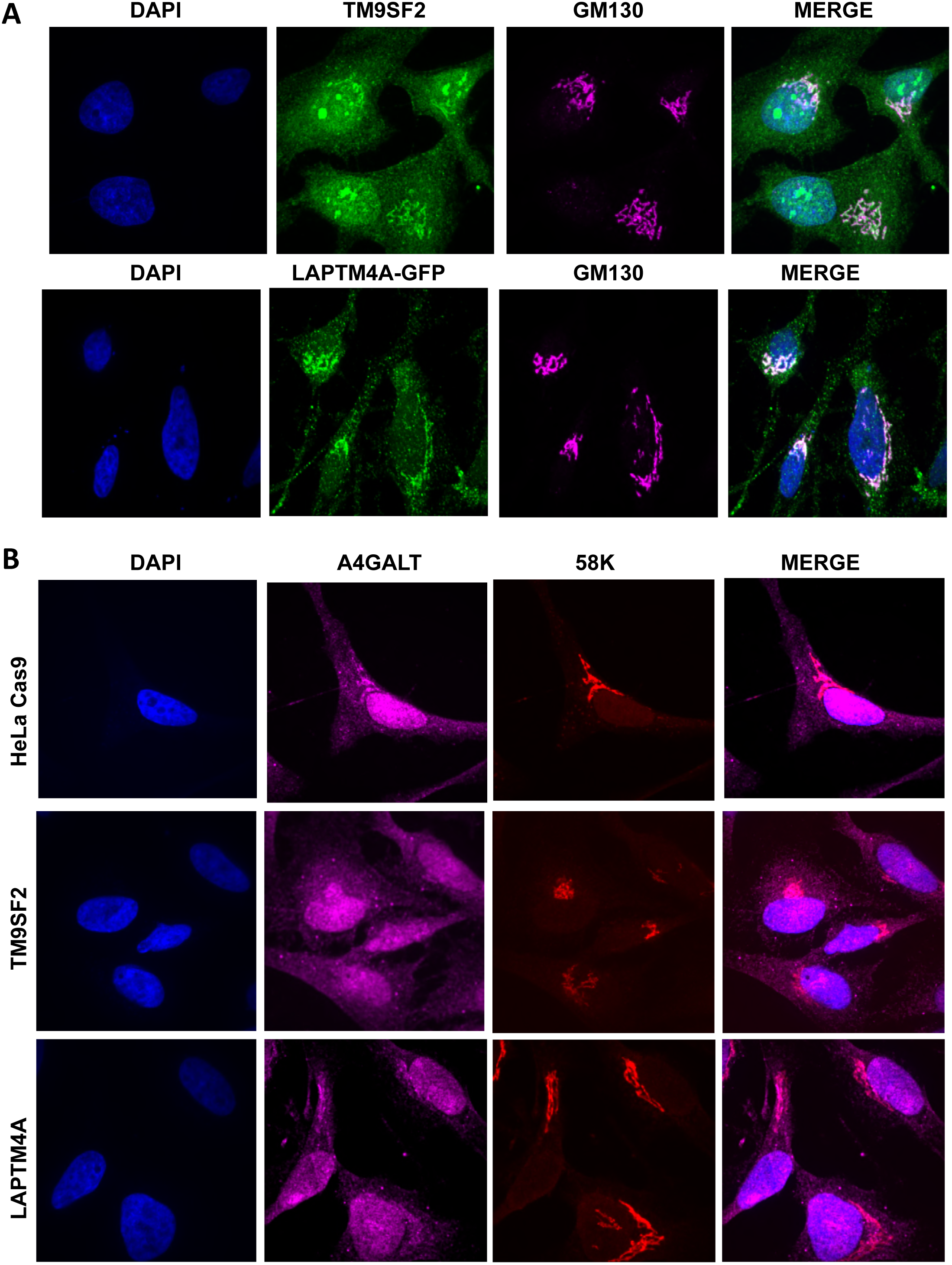
Subcellular localization of TM9SF2, LAPTM4A and A4GALT in wt and mutant HeLa cells. A). Confocal immunofluorescence microscopy of HeLa cells stained with anti-TM9SF2 (A; green) anti-GM130 to label the Golgi (pink)and DAPI. For LAPTM4A localization, HeLa cells were transfected with GFP-tagged LAPTM4A which was imaged directly after counterstaining as above. B) Confocal immunofluorescence microscopy of control and mutant HeLa Cas9 cells labeled with anti-A4GALT antibody (pink), anti-58K (red; labels Golgi) and DAPI (blue, labels nuclei). GM130 and 58K stain similar populations of Golgi membranes and were used interchangeably to accommodate the primary antibodies of interest.

The subcellular distribution of TM9SF2 and LAPTM4A raised the possibility that these proteins might enable EHEC infection by facilitating Gb3 biosynthesis, either by participating in Golgi-localization of precursor substrates or enzymes specifically, or by acting more generally as matrix proteins to ensure overall Golgi integrity. To investigate the former possibility, we immunolabeled the A4GALT enzyme in wt and in TM9SF2 and LAPTM4A cells. A4GALT maintained proper Golgi localization in both mutant cell lines (Fig. 5B), suggesting that TM9SF2 and LAPTM4A are not required either for the production or distribution of A4GALT. To investigate if TM9SF2 and LAPTM4A might instead facilitate Gb3 trafficking by acting more generally in Golgi integrity, we performed qualitative and quantitative image analysis of both cis-medial- and trans-Golgi compartments. GM130 (a cis-medial Golgi marker) appeared morphologically normal in the mutant cells (Fig. S5A) and quantitative characterization of the trans-Golgi, where the final step in Gb3 biosynthesis is thought to occur, did not reveal differences in either the integrity (as measured by confocal Z-stack nominal 2- dimensional area) or localization (as measured by nuclear centroid displacement) of this sub-compartment (Fig. S5BC). Further analyses will be required to identify the precise defect that leads to glycosphingolipid deficiency in these mutants.

## Discussion

EHEC encodes two potent virulence factors that empower it to disrupt the colonic epithelium during infection: its T3SS, which enables intimate attachment of bacteria as well as translocation of multiple effectors that disrupt epithelial cell processes, and Stx, a potent translation inhibitor that triggers multiple stress responses in cells within and outside of the intestinal tract. These virulence factors were acquired by horizontal transmission in distinct steps in the pathogen’s evolution (54) and are generally thought of as functionally independent. However, our CRISPR/Cas9-based screen for host mutants with reduced susceptibility to EHEC infection uncovered a remarkable overlap in host factors that mediate the response to these bacterial products. The screen for mutations enriched after infection with Stx+ and T3SS+ EHEC identified numerous loci known to be associated with sphingolipid and glycosphingolipid biosynthesis, in particular factors required for production of the Stx receptor Gb3, as well as two loci (TM9SF2 and LAPTM4A) with largely undefined cellular roles that were also required for toxin binding. Unexpectedly, mutants lacking these factors are also less susceptible to cytotoxicity associated with EHECs T3SS. These mutations interfered with early events associated with T3SS and Stx pathogenicity, markedly reducing entry of T3SS effectors into host cells and binding of Stx. Although the means by which these host loci and the processes associated with them are exploited by EHEC are not fully understood, the convergence of Stx and T3SS onto overlapping targets raises intriguing possibilities for design of therapeutic agents countering EHEC infection.

Previous studies of Stx and EHEC T3SS have characterized some of the pathways through which these factors act upon host cells, such as the binding and retrograde transport that enables Stx to reach its intracellular target, the stress responses induced by Stx, the processes underlying formation of pedestals, and the targets and mechanisms of effectors (55–58). Notably, our findings suggest that disruption of only a subset of host genes provides protection against cytotoxicity when both virulence factors are present. Interestingly, we identified loci that influence early steps within virulence pathways, e.g., loci that are required for Stx binding to host cells or T3SS effector translocation rather than loci that mediate toxin trafficking (e.g., clathrin, dynamin, SNX1/2) (18) or interact with translocated effectors (e.g., N-WASP, IRTKS, IRSp53) (58–60). Such proximal factors have also been identified in other genome-wide CRISPR screens for host mutants resistant to cytotoxicity by other pathogens. For example, a Yersinia RNAi screen and our *Vibrio parahaemolyticus* CRISPR screen yielded loci that reduced effector translocation (23, 45). For EHEC virulence factors, disrupting early steps in their interactions with host cells may be particularly protective because these virulence factors disrupt multiple cellular processes once internalized; for example, inactivation of the host response to a single T3SS effector may still leave cells vulnerable to the activity of other growth-interfering effectors.

Early steps in the interactions between host cells and EHEC’s virulence factors were also likely identified in our screen because of the previously unrecognized overlap between host factors utilized by T3SS and Stx at the start of their encounters with epithelial cells. We found that mutants with disrupted synthesis of Gb3, which were expected to be resistant to Stx-mediated growth inhibition, also exhibited an unexpected reduction in their sensitivity to T3SS-mediated cytotoxicity that was associated with reduced translocation of Tir. A majority of these mutants also exhibited reduced susceptibility to EPEC infection, suggesting that common host processes may mediate the actions of EHEC and EPEC’s related T3SS.

Although many mutants identified by our screen share the characteristic of lacking Gb3, it is unlikely that this deficit is the sole factor underlying their resistance to T3SS. Mutants lacking A4GALT, UGCG, TM9SF2 and LAPTM4A appear equally devoid of extracellular Gb3 in assays of Stx binding; however, they exhibit varying degrees of resistance to Δ*espZ* EHEC. T3SS resistance is also not fully correlated with the extent to which Tir translocation into these cells is reduced. These observations suggest that the reduction in Gb3 levels is associated with additional cellular changes (e.g., in overall sphingolipid homeostasis, membrane/lipid raft composition, or intracellular trafficking) that also modulate the host response to EHEC infection.

The means by which mutations in TM9SF2 and LAPTM4A prevent accumulation of Gb3 remain to be determined. We found that both proteins are localized within the Golgi, raising the possibility that they modulate the activity, localization, or transport of glycosphingolipid biosynthetic factors, which also occurs within this organelle. TM9SF2 was previously found to regulate the localization of NDST1, a Golgi localized enzyme that catalyzes N-sulfation of heparan sulfate, and to be required for accumulation of NDST1’s reaction product (28). TM9SF2 and the associated heparan sulfate N-sulfation are important for host cell binding and entry by CHIKV virus (28); however, the absence of other hits associated with heparan sulfate in our screen suggests that this phenotype is not related to our results. Minor abnormalities in several other glycosylation pathways were also associated with TM9SF2 disruption, but the underlying mechanism was not determined. We found that TM9SF2 is not required for correct localization or accumulation of A4GALT, suggesting that TM9SF2 may act prior to the terminal step of Gb3 synthesis. Similarly, LAPTM4A mutation did not appear to modulate A4GALT production or localization. It is unclear why previous studies have observed LAPTM4A in lysosomes and late endosomes rather than the Golgi localization that we detected; further studies will be needed to dissect the targeting and activity of LAPTM4A and its relationship to production of Gb3. Protein annotation and studies in the mouse homolog (MTP) suggest LAPTM4A may be involved in intracellular transport of nucleosides (61). Together with its Golgi localization shown herein, LAPTM4A could be involved in transporting activated sugars to the Golgi lumen, to supply precursors for Gb3 biosynthesis.

Although EHEC is susceptible to common antibiotics, antibiotic treatment is generally contraindicated during EHEC infection, as antibiotics can increase production and release of Stx, leading to the development of HUS (62). A variety of alternative therapies have been proposed to counter the effects of toxin, including compounds that sequester or neutralize toxin, block its binding to host cells, or disrupt toxin internalization, processing, or intracellular activity (63). Their activity has largely been tested in toxin-treated cell lines; a few have also been studied in animal models, but not in the context of EHEC infection. Our results suggest that a subset of these compounds, namely those that alter production of Gb3, may reduce pathogenesis associated with T3SS as well as Stx, and thus may be particularly effective in countering EHEC infection. Further studies of these and related compounds may enable identification of agents that counter host susceptibility to translocation of EHEC’s T3SS effectors as well as to the effects of Stx, which could hold high therapeutic potential. Thus, our identification of host factors related to both T3SS and Stx susceptibility provides guidance in prioritizing the development of therapeutics aimed at countering EHEC pathogenesis.

## Materials and Methods

### Bacterial strains, plasmids and growth conditions

All bacterial strains and plasmids used in this study are listed in Table S1. Primers used in strain construction are shown in Table S1. Bacterial strains were cultured in LB medium or on LB agar plates at 37°C unless otherwise specified. Antibiotics and supplements were used at the following concentrations: carbenicillin: 50μg/ml; ampicillin: 100ug/mL; chloramphenicol: 20μg/ml; kanamycin: 50μg/ml; streptomycin: 200μg /ml; IPTG: 1ug/ml. The EHEC Δ*espZ* deletion mutant was constructed by allelic exchange using a derivative of the suicide vector pDM4 that included *espZ*-flanking sequences (64). Deletion of *stx1*, *stx2* and *escN* was performed using lambda-red mediated recombination (65).

### Eukaryotic cell lines and growth conditions

HT-29, HeLa and 293T cells and their derivatives were cultured in DMEM supplemented with 10% fetal bovine serum. Cells were grown at 37°C with 5% CO2 and routinely passaged at 70-80% confluency; media was replenished every 2-3 days.

### Positive selection screen using the HT-29 CRISPR AVANA libraries

The HT-29 libraries were constructed as described (23) using the AVANA sgRNA library, which contains four sgRNAs targeting each of the human protein coding genes (26). For each library, two sets of 7 T225 flasks were seeded with 12.5*10^6^ cells per flask, then incubated for 48 hours. At the time of the screen there were 175*10^6^ cells per experimental condition, corresponding to ~2000X coverage per perturbation. Cells were at ~70% confluency at the time of infection. One set of flasks served as an uninfected control; the second was infected with the EHEC Δ*espZ* strain. Cells were harvested from the control at the time of infection.

For infection, the Δ*espZ* strain was grown overnight statically in LB media to an OD_600nm_ of 0.6, then centrifuged and resuspended at OD_600nm_ 0.5 in DMEM. HT-29 cells were infected with EHEC Δ*espZ* at an MOI=100 and incubated under standard culture conditions for 6 hours, with a media change at 3 hr post-infection to remove nonadherent bacteria and prevent media acidification. At 6-hour post-infection, cells were washed 3 times with 1X DPBS to remove nonadherent bacteria, then replenished with fresh media supplemented with 1X antibiotic-antimycotic solution (ABAM) (Gibco-containing 100ug/mL streptomycin) and gentamicin 100ug/ml (Gent) (“stop medium”). After overnight incubation, fresh “stop medium” was added to each flask. Flasks were monitored daily by inverted light microscopy to follow the recovery of survivor cells. Upon reaching 70% confluency, the cells were trypsinized, pooled and re-seeded for the next round of infection, always keeping a minimum number of 80*10^6^ cells to maintain a coverage of at least 1000X. The infection and selection procedure was repeated for the second, third and fourth rounds of infection. Additionally, a subset of cells from each population were used for preparation of genomic DNA.

### Genomic DNA preparation, sequencing and STARS analyses of screen results

Genomic DNA (gDNA) was extracted from 100×10^6^ input cells (uninfected) and after each round of infection with EHEC Δ*espZ* (rounds 1, 2, 3 and 4) using the Blood and Cell Culture DNA Maxi Kit from Qiagen. The gDNA was subjected to PCR to amplify guide RNA sequences as previously described (26). The read counts were first normalized to reads per million within each condition by the following formula: reads per sgRNA/total reads per condition × 10^6^. Reads per million were then log2-transformed by first adding 1 to all values, in order to take the log of sgRNAs with zero reads. For analyses, the log2 fold-change of each sgRNA was determined relative to the input sample for each biological replicate (Table S2). The STARS algorithm for CRISPR-based genetic perturbation screens was used to evaluate the rank and statistical significance of the candidate genes as described (26).

### Construction of HT-29 Cas9 and HeLa cells with targeted gene disruptions

The sgRNA sequences used for construction of HT-29 Cas9 mutant cells are shown in Table S3. All sgRNA oligo sequences were obtained from Integrated DNA Technologies and cloned into the pLentiGuide-Puro plasmid as previously described (23). Briefly, 5μg of plasmid pLentiGuide-Puro was digested with *BmsBI* (Fermentas) and purified using the QIAquick Gel extraction kit. Each pair of oligos was annealed and phosphorylated with T4 PNK (NEB) in the presence of 10X T4 DNA ligase buffer in a thermocycler with the following parameters: i) incubation for 30 minutes at 37°C, ii) incubation at 95°C for 5 min with a ramp down to 25°C at 5°C per minute. Oligos were then diluted 1:200 and 1μl of the diluted oligo mixture was ligated with 50ng of *BsmBI* digested plasmid. Ligations were transformed into STBL3 bacteria, and transformed clones were checked by PCR and DNA sequencing. sgRNAs cloned into pLentiGuide-Puro were transduced into HT-29 Cas9 cells as described below, and after 10 days of selection with Puromycin (1μg/ml), the extent of disruption of the targeted gene was analyzed by immunoblotting for the corresponding gene product or Sanger sequencing (Fig. S2C).

### Lentivirus Preparation and Transductions

Lentiviral transductions were performed as previously described (23). Briefly, all lentiviruses were made by transfecting 293T cells using TransIT-LT1 transfection reagent, the lentiviral packaging plasmids psPAX2 and pCMV-VSVG and the corresponding cargo plasmid according to the manufacturer’s protocol. 48h following transfection, 293T culture supernatant was harvested, filtered through a 0.22μm pore filter, and added to target HT-29 or HeLa cells grown to 70-80% confluency in 6-well plates; a second virus supernatant were harvested 72 hr after transfection and added to target cells. After each virus’ supernatant addition to HT-29 cells, spin infection was performed by adding 8μg/ml polybrene and spinning the 6-well plates were at 1600g for 2h at 30°C; HT-29 cells were then returned to 37°C. Puromycin selection for positive transductants was initiated the following day. For transduction of HeLa cells, spin infection was not performed.

### Cell survival assays

For cell survival assays, 5×10^5^ HT-29 cells were seeded into 6-well plates and grown for 48 hours in DMEM supplemented with 10% FBS. EHEC (or EPEC) strains for infections were prepared as for library infections described above. HT-29 cells were infected at an MOI=100 (or with uninoculated media in the case of mock infection), with media changes and infection termination as for library infection. Mock infected cells were fed but not passaged during the outgrowth period. Following infection and outgrowth for 1 or 5 days, cells were quantified by Trypan Blue (0.4%trypan blue) exclusion using a Countess II Automated Cell Counter (Thermo Fisher Scientific). Cell survival after EPEC infection was measured 4 hr post-infection.

### Stx cytotoxicity assay and measurement of Stx released during infection

HT-29 cells were seeded at 1×10^6^ cells/ well the day before the assay. Cell monolayers were then exposed to a range of concentrations of pure Stx1 or Stx2 holotoxins for 6 hours. Cell survival was measured by Trypan Blue exclusion as described above, then % of survival was calculated in comparison to HT-29 controls that did not receive toxin treatment. Stx released during infection of HT-29 cells at 3hr and 6hr was measured by ELISA, as described (66).

### Tir Translocation Assays

The Tir translocation assay was performed as previously described (47). Briefly: HT-29 cells were plated at 1×10^5^ cells/well in 96 wells and assayed at confluency. EHEC strains harboring Tir-CyaA fusion or CyaA vector control were grown in LB overnight then diluted 1:100 in DMEM and grown to OD600 = 0.6 shaking at 37C. Cells were infected with MOI 100:1 for 90 minutes; cAMP was measured by ELISA using Biotrack cAMP kit (Amersham) according to manufacturer’s instructions.

### Immunoblot analyses

Mammalian cell lysates were prepared with RIPA buffer and protein concentrations were determined using BCA protein assay. 10ug of protein lysate was mixed with NuPAGE LDS sample buffer (Invitrogen) with 50mM DTT, separated by NuPAGE Bis-Tris gel electrophoresis and transferred to nitrocellulose membranes. Antibodies and concentrations used are listed in KEY RESOURCES TABLE. Blots were developed with the SuperSignal West Pico ECL kit, and imaging was performed on the Chemidoc Touch Imaging System (Biorad).

### Immunofluorescence

HT-29 cells or HeLa cells were seeded in 12-well plates on 18mm glass coverslips or 4- well chambers (Mat-TEK). Cells were fixed with 2% paraformaldehyde (PFA) for 20 minutes at room temperature, washed with 1X PBS 3 times, then permeabilized with 0.1% Triton X-100 in PBS for 30 minutes (except for cells stained for TM9SF2, which were subjected to combined fixation and permeabilization in ice cold methanol for 10 minutes). Cells were blocked in 5% normal goat serum in PBS (blocking buffer) for 1 hour, followed by overnight incubation with primary antibodies (Table S4) at 4°C. Cells were then washed 3 times with PBS followed by incubation with fluorescently-labeled secondary antibody for 1 hour at room temperature. Cells were counterstained with Alexa-568 Phalloidin and DAPI for actin cytoskeleton and nuclei, respectively. For extracellular binding of Alexa-488-tagged Stx, cells were not permeabilized; for intracellular binding, cells were permeabilized and stained as described above.

### LAPTM4A subcellular localization

HeLa Cas9 cells were plated on coverslip and transfected with LAPTM4A-GFP (Origene) per the manufacturer’s protocol (Mirus). 24 hours later, cells were processed for immunofluorescence as above. GFP was imaged directly without additional signal amplification.

### Fluorescent actin staining (FAS)

FAS assays were performed as described (67) with minor modifications. Briefly, HT29 cells were seeded at 1×10^6^ cell/well in 4-well chambers in DMEM + FBS. Three days after confluency, cells were infected with EHEC strains expressing GFP at an MOI=100 for 6 hours, with a media change after 3hr. After infection, cells were washed three times with PBS, fixed with 2% PFA and permeabilized with 0.2% Triton X-100. Cells were then stained with Alexa-633 Phalloidin and DAPI for visualization of actin cytoskeleton and cell nuclei. Slides were mounted using Prolong Diamond Antifade and analyzed by confocal microscopy. The experiment was repeated at least 3 times, and 250 cells were counted in total. The percentage of infected cells was determined by analyzing at least 20 random fields across different experiments; numbers of pedestals were determined by counting AE lesion in 100 infected cells. All comparisons were relative to HT-29 cells.

### Lipid raft assay

For imaging of GFP-GPI, the indicated cells were split into 12-well glass bottom plates (MatTek). 1 day later, cells were transfected with GFP-GPI using TransIT-LT1 reagent following the manufacturer’s recommended protocol (Mirus). 24 hours later, cells were washed and then imaged in FluoroBrite-DMEM (Invitrogen), with live fields of single confocal slices of cell bottoms taken for 1 minute using 1 second exposures at 75% laser power. ROI mean intensities (with ROI drawn to avoid overlapping cell protrusions and saturated pixels) were calculated for each frame using the Plot Z-axis profile function in ImageJ. Greater than 20 cells and at least 20 movies were analyzed for each condition. Results are expressed as mean +/- SEM.

### Golgi complex morphological analyses

Golgi integrity was assayed by calculation of the mean distance between the manually defined weighted centroid of nucleus (as defined by DAPI staining) and trans-Golgi (as defined by TGN46 staining), and from 2-dimensional area of manually defined ROI of maximum-intensity projection of individual confocal slices of TGN46 staining. At least 20 cells were analyzed for each condition. Results are expressed as mean +-SEM.

### Shiga toxin labeling and Flow Cytometry (FACS)

Shiga toxins 1 and 2 (holotoxins) were obtained from Tufts Medical Center; cholera toxin was purchased from Sigma. All toxins were diluted in PBS and labeled with Alexa488 or Alexa647 micro-labeling kit (Invitrogen) according to manufacturer’s instructions. For FACS analysis of Stx binding, HT-29 cells were seeded at 5×10^5^ cells/well in 6-well plates, while HeLa cells were seeded 2.5×10^5^ cells/well, then incubated for 24 hours. Cell monolayers were washed 3 times with EBSS, trypsinized, resuspended in PBS with labeled Stx (10nM) or CT (1nM), and incubated on ice for 30 min. Cells were then centrifuged, resuspended in FACS buffer (DPBS + 10% FBS) and analyzed by flow cytometry.

### Statistical Methods

Statistical analyses were carried out using a one-way ANOVA with Dunnet post-correction on GraphPad Prism5.

## Acknowledgements

We gratefully acknowledge Professor James Kaper (University of Maryland) for providing EPEC strains and the Tir-cyaA plasmid, to Professor Yusuke Maeda (Osaka University) for providing anti-TM9SF2 antibody and to Professor Peter Howley for HeLa Cas9 parental cells. We thank Waldor lab members for helpful discussion about this project. This work was supported by HHMI and R37 AI-042347 (MKW), B.S. was supported by National Sciences and Engineering Research Council of Canada (NSERC) PGS-D award (487259), J.E.L. was supported by the NIH under NRSA T32AI007061 from the NIAID.

---

**Figure S1. Cytoxicity and host cell survival associated with various EHEC strains and purified toxin.** A) Graphs show the abundance of HT29 cells infected with the indicated strain relative to the abundance of mock-infected cells at day 1 post infection with EHEC strains. Data reflect the mean +/- SD (n=3). P values (* P<0.05, ** P<0.01, **** P<0.0001) **B)** Kinetics of HT-29 cell death and recovery after challenge with Δ*espZ* (red), the Shiga toxin-deficientΔ*espZ* Δ*stx1* Δ*stx2* (blue), the T3SS-deficient ΔespZ ΔescN (green), or mock infected. Data are representative of 3 independent experiments. C) Abundance of Shiga toxins 1 and 2 in media during infection of HT-29 cells with Δ*espZ* and Δ*espZ*Δ*stx1*Δ*stx2* (negative control). Toxin levels were assayed at 3 hr post infection and at 6 hr post infection (3 hr post media change), using an ELISA with antibody 4D1 which detects both toxins. D) Survival of HT-29 cells after 6 hours intoxication with either pure Shiga toxin 1 or 2; UD = undetectable.

**Figure S2. CRISPR screen results and validation of mutations generated in candidate loci**. A) Box plots showing the distribution of sgRNA frequencies in each HT-29 CRISPR library prior to infection and following each round of infection with Δ*espZ* EHEC. Line in the middle of the box indicates the median and whiskers comprise the 5th to 95th percentile. B) Heatmap of sgRNA enrichment in each HT-29 CRISPR library after successive rounds of Δ*espZ* EHEC infection. The heatmap shows each of the 4 sgRNAs targeting the genes; the darkness of the blue color correlates with the fold-enrichment of the sgRNA compared to the input libraries. **C)** Western blot of whole cell lysates of HT-29 Cas9 cells and CRISPR mutants. Arrows indicate the molecular weight corresponding to each target protein. Antibodies used for validation are listed in Table S4. **D**) Analysis of indels in HT-29 mutants. Trace files show sequence reads indicating gene disruption at the sgRNA binding site on A4GAL and LAPTM4A mutants, compared to the gene in the parental cell line (WT). Red box outlines the sgRNA sequence.

**Figure S3.** A) Single channel and merged images corresponding to merged images shown in Fig 3C generated from confocal microscopy of control and mutant HT-29 Cas9 cells infected for 6 hr with GFP-producing EHEC, then stained with Alexa-647-phalloidin and DAPI. Arrows in merged images indicate pedestals (arrow). B) Graphs show the abundance of HT29 cells infected with the indicated EPEC strain relative to the abundance of mock-infected cells 4hr-post-infection with EPEC. Data reflect the mean +/- SD (n=3). P values (* P<0.05, ** P<0.01, # P<0.0001) C) Abundance of control and mutant HT29 Cas9 cells infected with *espZ* and *escN* EPEC relative to the abundance of mock-infected cells at 4 hr post-infection. Data correspond to mean and SD from 3 independent experiments. P values (** P<0.01) D) Analysis of lipid rafts components in control and mutant HeLa cells. Representative confocal slice of adherent cell bottom, 24 hours after transfection with GFP-GPI which traffics to the plasma membrane and inserts preferentially into lipid rafts. E) Quantitation of lipid rafts in control HeLa Cas9 cells and mutants. Total plasma membrane fluorescence (arbitrary fluorescence units) is depicted, along with kinetics of fluorescence decay with quantitative photobleaching. Data represent mean/SEM.

**Figure S4. Flow cytometry analyses of toxin binding to control and mutant host cells.** A) Flow cytometry analysis of Stx2-Alexa647 binding to control and mutant HeLa Cas9 cells. Histograms show HeLa cell population in the presence (pink) or absence (green) of toxin. B) Flow cytometry analysis of CT-Alexa647 binding to control and mutant HT-29 cells Histograms show HT-29 cell population in the presence (pink) and absence (green) of toxin.

**Figure S5. Visualization and quantitative analysis of Golgi structure in control and mutant host cells.** A) Confocal immunofluorescence microscopy of Golgi structure in control and mutant Hela Cas9 cells. Cis-medial Golgi (pink) were stained with anti-GM130, and nuclei (blue) were stained with DAPI. B) Confocal immunofluorescence microscopy of the trans-Golgi network (TGN) in control and mutant Hela Cas9 cells. TGN was stained with TGN46 (green) and nuclei (blue) were stained with DAPI. C) Quantitative analysis of TGN morphology in control and mutant Hela Cas9 cells, based on cell staining shown in (B). Distance of TGN from nuclei (left) and nominal TGN area (right) were determined for at least 20 cells.

**Table S1. Primers used to construct EHEC EDL933 mutants**

**Table S2. STARS analysis of Library B round 4**

**Table S3. Sequence of sgRNAs and plasmids used to construct HT-29 Cas9 and HeLa Cas9 CRISPR mutants**

**Table S4. Antibodies used for western blot and immunofluorescence**

